# The Bayesian polyvertex score (PVS-B): a whole-brain phenotypic prediction framework for neuroimaging studies

**DOI:** 10.1101/813915

**Authors:** Weiqi Zhao, Clare E. Palmer, Wesley Thompson, Terry L. Jernigan, Anders M. Dale, Chun Chieh Fan

## Abstract

The traditional brain mapping approach has greatly advanced our understanding of the localized effect of the brain on behavior. However, the statistically significant brain regions identified by standard mass univariate models only explain minimal variance in behavior despite increased sample sizes and statistical power. This is potentially due to the generalizable explanatory signal in the brain being non-sparse, therefore not captured by the thresholded, localized model. Here we introduced the Bayesian polyvertex score (PVS-B), a whole-brain prediction framework that aggregates the effect sizes across all vertices to predict individual variability in behavior. The PVS-B estimates the posterior mean effect size at each vertex with mass univariate summary statistics and the correlation structure of the imaging phenotype, and weights the imaging phenotype of participants from an independent sample with these posterior mean effect sizes to estimate the generalizable effect of a brain-behavior association. Empirical data showed that the PVS-B was able to double the variance explained in general cognitive ability by an n-back fMRI contrast when compared to prediction models based on the mass univariate parameter estimates as well as models in which only vertices thresholded based on p-value were included. A fivefold improvement in variance explained by the PVS-B was observed using a stop signal task fMRI contrast to predict individual variability in the stop signal reaction time. We believe that the PVS-B can shed light on the multivariate investigation of brain-behavioral associations and will empower small scale neuroimaging studies with more reliable and accurate effect size estimates.

## INTRODUCTION

In traditional neuroimaging analyses, a brain-behavior association is assessed with a brain mapping approach where the associative effect on behavior is estimated independently at each measured unit of the brain data, e.g. vertex or voxel. These mass univariate effects are subsequently summarized as a parametric map where statistical correction methods are applied to map out a set of brain regions with significant behavioral associations. Given the limited statistical power (Cremers, Wager & Yarkoni, 2017) and signal-to-noise ratio (SNR; Gonzalez-Castillo et al., 2012) of neuroimaging studies, a brain-behavior relationship is usually localized to a few clusters of brain regions with the most significant p-values. With growing numbers of imaging consortia and increased collaborative effort for data sharing, the sample size of neuroimaging studies is large enough to detect small brain-behavior relationships spanning a larger number of vertices or voxels; however, these mass univariate effects individually explain a very small percentage of between subject variability in behavior (Poldrack et al., 2017). Large effect sizes are rarely observed at the brain region or vertex level (Stanfield et al., 2008). With a sample size of more than 14,000 participants, Smith and Nichols (2017) demonstrated that a statistically significant imaging composite measure explained less than 1% of the variance in behavior even after Bonferroni correction of 14 million tests. This questions whether those small effect sizes detected by large scale imaging studies are really meaningful or generalizable. Indeed, neural signatures distributed across the cortex, spanning multiple sub-networks, have been shown to perform better than specific, predefined brain regions or networks at classifying clinical disorders (Bruin, Denys, & Wingen, 2019; Reddan, Lindquist, & Wager, 2017) and predicting individual differences in cognitive processes (Chang et al., 2015). The predictive power of a given brain phenotype is not localized but appears to be widespread across the cortex.

Similar observations were made in the field of genetics. Genome-wide association studies (GWAS) use mass univariate regressions across the genome to localize genetic loci associated with behavior in thousands of participants. However, the detected genetic loci that survive Bonferroni correction often only account for a fraction of the variance in complex human phenotypes. As a test to investigate the magnitude of generalizable signals among those non-significant small effect genetic variants, polygenic risk scores (PRS) were developed as an aggregated sum over all of the effect sizes from GWAS (Davies et al., 2011; Le Hellard & Steen, 2014; Torkamani et al., 2018; Yang et al., 2010). By pulling together the effects of many informative but not necessarily statistically significant SNP loci, the PRS explained a much higher proportion of behavioral variation than only the significant SNPs. When the true effects are less sparse, meaning the effects are small and ubiquitous across the genome, the PRS out-performs prediction models with variant sets selected based on statistical significance (Dudbridge et al., 2013). In particular, the invention of the PRS has enabled researchers of smaller scale studies to make powerful inference using the effect size estimates from large-scale GWAS studies (Torkamani et al., 2018).

Given the observed similarity across fields, we explored that whether we could adapt the traditional brain mapping approach to a framework for prediction using whole-brain phenotypes. We developed a novel imaging analysis framework: the polyvertex score (PVS). The PVS harnesses the explanatory power of an imaging phenotype by incorporating information across all voxels or vertices that are readily available from the brain mapping analysis. It aggregates the effect sizes across all vertices of the cortical surface for behavioral prediction, mirroring the function of PRS. Two versions of the PVS were developed. The mass univariate PVS (PVS-U) is a summary measure of all the univariate linear associations across the cortex. The Bayesian PVS (PVS-B), on the other hand, uses a Bayesian parameter estimation process that incorporates the correlation structure, estimated SNR, sample size, and number of vertices of the brain phenotype. This accounts for the non-independence between vertices in the brain, which is a limitation of the mass univariate approach.

Using simulations and functional neuroimaging data from the Adolescent Brain Cognitive Development (ABCD) study, we tested two main hypotheses. Firstly, we hypothesized that using all estimated mass univariate effects across the cortex, regardless of statistical significance, would better predict individual variability in behavior compared to predictions based on a subset of the most significant vertices. This would support the observation that the explanatory effect of cognitive behaviors is widespread across the cortex and that vertices below the traditional, corrected threshold are informative for brain-behavior relationships. Secondly, we hypothesized that incorporating the estimated SNR of the imaging phenotype and the correlation structure across vertices would improve our predictive power; thus, prediction models based on the Bayesian parameter estimates of brain-behavior associations (PVS-B) would explain more individual variability compared to PVS-U.

## METHOD

### A background based on the brain mapping approach

Traditionally, the association between an imaging phenotype and a behavior is tested with univariate regression models at each vertex. The effect size of a brain phenotype on behavior, based on the brain mapping approach, estimates a linear, additive relationship between the imaging measurement, *X*, at each vertex, and the phenotypic outcome, *y*, as:

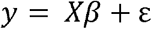

for *X* denotes a standardized *N* by *V* matrix of a given imaging phenotype where *N* and *V* denotes the number of subjects and vertices respectively. *y* represents an *N* by 1 vector of a behavioral phenotype, and *β* is the *V* by 1 vector of parameters of interest. With the goal of localizing the effect rather than estimating the total explanatory signal, the brain mapping approach omits the correlation information among vertices to reduce the computational demand.

Recent debates on the reproducibility and the small effect sizes of neuroimaging research are based on such mass univariate estimates from the brain mapping framework (Poldrack et al., 2017; Smith and Nichols, 2017). We argue that inferring the magnitude of a brain-behavior association based on these mass univariate estimates requires further consideration. Generalizing based on the effect size estimates of the most significant vertices/ROIs of the mass univariate model commits the assumption that the underlying true signal is sparse and localized, and clusters of vertices/ROIs with minimum P values (Min-p) contain the main source of generalizable signals. However, as previously mentioned, the explanatory power of the brain on behavior appears to be nonsparse, and thus cannot be captured by the most significant vertices/ROIs. In order to generalize the effect sizes of the whole brain phenotype, we need a prediction framework that accounts for the nonsparseness of the brain signal on behavior.

Rooted in this brain mapping approach, we proposed the PVS estimation and prediction framework. Importantly, we designed the parameter estimation process to enable smaller scale imaging studies to make inferences based on the effect size estimates from large scale imaging consortiums, mirroring the contribution of the PRS to genetics research. Therefore, all estimation processes only require summary statistics, i.e. the effect size estimates of the mass univariate regression models of the brain mapping framework. There is no need for original individual level data if the summary statistics have been shared by the research community, such as the design of ENIGMA project (Stein et al., 2012).

### Mass univariate parameter estimation

In the brain mapping context, the relationship between all vertices of the brain and the behavior is estimated independently at each vertex with a univariate model. The effect of each column of the brain matrix, *X*, is estimated independently and the correlation matrix of the brain phenotype, *X*′*X*, is set to *I*, the identity matrix. The parameter estimates based on a mass univariate model is thus reduced to the form:

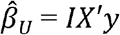

Independent estimation of the parameter estimate at each vertex effectively reduces the computational demand of the mass univariate model. However, ignoring the correlation structure among vertices could result in the biased estimation of the multivariate beta parameters, hampering the ability of mass univariate models to make accurate inferences and predictions. A parameter estimation approach that accounts for the correlation structure of the brain phenotype is thus called for.

### Bayesian parameter estimation

To tackle the correlated signal of the imaging phenotype at each vertex, we developed a Bayesian parameter estimation approach where the correlation information across vertices is incorporated into the parameter estimation process. Similar framework has been proposed in the field of genetics (Vilhjálmsson et al., 2015). The intuition behind the formulation of the PVS-B is to approximate the multivariate linear regression coefficients given the massive univariate beta estimates while providing analytically derived empirical priors to ensure the information matrix is positive definite.

The Bayesian parameter estimation procedure calculates the posterior mean effect sizes of the brain phenotype by weighting the mass univariate beta estimates with a factor that accounts for the observed correlation structure of the cortex and the experimental condition of the imaging data:

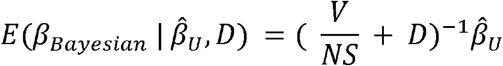

Here, the posterior mean effect sizes 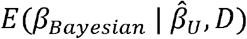, are approximated by the mass univariate beta estimates 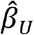 multiplied by the inverse of the correlation structure of the brain, *D*, and a shrinkage factor that accounts for the number of vertices, *V*, the number of participants, *N*, and the SNR of the brain-behavior association, *S*.

The SNR of the brain phenotype, *S*, is estimated using the moment estimator on the mean effects (Schwartzman et al., 2017), which characterizes the amount of signal given the observed associations. Using the mass univariate beta estimates, *S* can be estimated by:

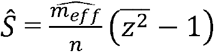

where 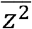 is the mean of the squared z-statistics of the mass univariate regressions across vertices, and the 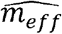 is the estimated effective number of vertices. 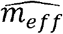 is calculated by the number of vertices, *V*, divided by the second spectral moment of the correlation matrix, *D*, a factor that captures the effect of the correlation structure of the brain.

The benefits of implementing this Bayesian parameter estimation are 2 folds: 1) the Bayesian parameter estimation procedure takes into account the correlation structure of the brain phenotype, circumventing the independence assumption of the mass univariate approach; and 2) the SNR of the brain phenotype is being incorporated. Given that imaging data usually have low SNR, accounting for the SNR in the parameter estimation process can reduce the variance in prediction error that leads to better predictive performance. We note that the PVS-B implemented in this study explicitly assumes a prior that all vertices have true associations on behavior.

### Behavioral prediction

#### Polyvertex scores

Motivated by the PRS analysis for GWAS, a polyvertex score (PVS) can be calculated from neuroimaging data by aggregating the explanatory power of all vertices on behavior. The PVS represents the predicted behavioral phenotype based on the cortex-wide associations between the imaging phenotype and the observed behavior. We implemented two types of PVS that utilize the mass univariate and Bayesian parameter estimates respectively. A mass univariate PVS (PVS-U), based on the mass univariate parameter estimates, was computed as the brain phenotype at each vertex for an individual multiplied by the mass univariate parameter estimates acquired from an independent sample:

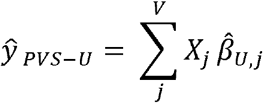

The PVS-U summarizes the effect size at all vertices on individual variability in behavior, with the assumption of independence at each vertex.

Similarly, a Bayesian PVS (PVS-B) was calculated using the Bayesian parameter estimates:

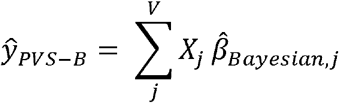

The PVS-B is hypothesized to harness the multivariate effect of an imaging phenotype on behavior by accounting for the correlation structure and the SNR of the brain phenotype, and should yield superior predictive performance over the PVS-U.

#### Thresholding

Modeling a brain-behavior relationship with a whole-brain phenotype yields significant gain in prediction accuracy when the underlying true association between the brain and behavior is widespread across the cortex. However, when the true signal is sparse, a whole-brain phenotype risks inferior prediction performance compared to methods that include only the most significant vertices for prediction. In the current study, we did not know the sparsity of the true brain-behavior associations, therefore we implemented 3 levels of thresholding for the PVS-U and PVS-B. Specifically, we tested whether thresholding the number of vertices included in the PVS-U and the PVS-B improved prediction performance when the signal sparsity was high. The thresholding procedure was performed as follows: we ranked the absolute effect sizes for all vertices and removed those ranked lower than a threshold proportion. Three levels of thresholding were implemented such that the top 50%, 10% and 1% of vertices were retained for the PVS-U or PVS-B.

To link our predictive methods with the canonical statistical inference approach where a brain and behavior relationship is established when any single vertex shows a significant association with the behavior, we compared our methods with the predictive performance of the vertex with the most significant mass univariate z-score which we have referred to as the Min-p model.

We used simulations and empirically collected functional MRI data to examine prediction accuracy of the above mentioned 9 methods: the PVS-U, PVS-U 50% (PVS-U with 50% most significant vertices), PVS-U 10%, PVS-U 1%, the PVS-B, PVS-B 50%, PVS-B 10%, PVS-B 1%, and Min-p.

### Evaluating the performance of PVS

We used 10-fold cross validation to evaluate the generalization performance of PVS. The same training-testing schema was applied to each fold to ensure the independency between estimation and prediction. At each fold, mass univariate beta estimates obtained from the training set, containing 90% of the full sample, were multiplied by the imaging phenotype of each test set participant to obtain a PVS-U score (the predicted behavioral phenotype) for each participant in the test set. This procedure was repeated 10 times yielding a predicted behavioral score for each participant in the full sample. For the PVS-B, the posterior effect sizes were calculated with the estimated SNR, the correlation structure, and the mass univariate beta estimates from the training data, and multiplied by the imaging phenotype of each test set participant. Variance explained, *R*^2^, the squared correlation between the observed and predicted behavior phenotypes, was used as a metric for prediction accuracy.

### Simulations

First, we used simulations to assess this novel method and determine whether the different types of PVSs perform as expected in certain contexts. In particular, we assessed how signal sparsity, the SNR of the brain phenotype and sample size influenced the predictive accuracy of the above mentioned PVS methods. The proportion of the true signal, i.e. the signal sparsity level, was simulated at the levels of 100%, 50%, 10%, 1%, and 0.1% true signal. The 0.1% true signal level corresponds to the Min-p assumption where the significant effect lies in a single vertex. The SNR of the brain phenotype was simulated at the levels of 0.01, 0.05, 0.1, and 0.2. Each combination of signal and SNR was simulated independently 100 times, giving 2000 iterations in total.

For each iteration of the simulations, the predictive effect of the brain on behavior at each vertex, the true beta coefficient, was simulated as:

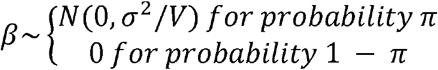

The beta coefficients were simulated by sampling independently from a standard normal distribution. A subset of these true beta coefficients was then set to zero as determined by 1 minus the proportion of true signal, *π*. For example, for an instantiation of 10% true signal scenario, 10% of the vertices were randomly assigned to have non-null effects which can account for *σ*^2^ of outcome variations in total, whereas the beta coefficients of the other 90% were set to zero.

Then, the simulated behavioral phenotype was calculated as a combination of the effect of an empirically collected brain phenotype, *X*, and independent noise weighted by the square root of the SNR of that iteration. The independent noise was sampled at the participant level from a standard normal distribution. To make the simulation more realistic, we used the empirical brain phenotype data as the independent variables, *X*, which was a 6103 by 1284 brain matrix of the 2back - 0back contrast of the nBack fMRI task of the baseline data of ABCD (ABCD Data Release 2.0.0; NDAR DOI:10.15154/1503209). *X* was smoothed at around FWHM 5mm, pre-residualized by age and categorical variables including sex, parent marital status, highest level of parental education, household income, self-reported race and ethnicity, and MRI scanner ID.

Within each iteration, two independent samples were randomly drawn at the sample size of 100, 500, 1000, 3000, 5000, and at the full sample size, 6103, to estimate the sample size dependency of prediction accuracy. Predicted behavioral phenotypes based on the PVS-U, the PVS-B, and their thresholded variants were calculated with 10-fold CV. Variance explained, *R*^2^, the squared Pearson correlation between predicted and simulated behaviors, served as a metric for predictive performance.

### Empirical Data

We examined the empirical utility of the PVSs by predicting individual variability of 2 different cognitive tasks with 2 fMRI contrasts using the baseline data of the ABCD Study (NDAR DOI: 10.15154/1503209). The ABCD Study is a longitudinal study across 21 data acquisition sites in the U.S. following 11,875 children starting at 9 and 10 years old. Detailed study designs and recruitment procedures, imaging acquisitions, and preprocessing pipelines are described in Garavan et al., (2018), B.J. Casey et al., (2018) and Hagler et al., (2018) respectively.

With the complete data of the ABCD study, we estimated the predictive performance of the vertex-wise *2 back - 0 back* contrast from the nBack fMRI task (Cohen et al., 2016) and the *correct stop vs. correct go* contrast from the Stop Signal Task (SST; Logan, 1994) on the *total composite score of cognition* from the NIH Toolbox (NIH-TB; Gershon et al., 2013) and the *stop signal reaction time* (SSRT) from the SST task respectively. Four brain-behavior associations of interest were examined: nBack predicting TC, nBack predicting SSRT, SST predicting TC, and SST predicting SSRT. All imaging and behavioral phenotypes were pre-residualized by age and categorical demographic variables including sex, parent marital status, highest level of parental education, household income, self-reported race and ethnicity, and MRI scanner ID. The final sample included 6103 participants for the nBack associations and 6472 participants for the SST associations.

Variance explained, *R*^2^, was calculated for each of the 9 prediction models for each association. To quantify the difference in predictive accuracy among methods, we calculated the bootstrap confidence interval (CI) for the variance explained with 1000 bootstrapped samples. These samples were generated by resampling with replacement the observed and predicted behavioral phenotypes based on the family structure of the complete sample of the nBack and SST fMRI tasks.

Lastly, to understand the effect of sample size on predictive performance, we calculated the out of sample variance explained *R*^2^ for each method at sample sizes of 100, 500, 1000, 2000, and 3000. The full sample was divided into 10 hold out samples based on family structures such that each holdout sample contained equal proportion of families and singletons. For each holdout sample, a thousand independent samples for each of the above-mentioned sample sizes were randomly selected from the remaining data. Methods were subsequently trained on these independent samples and generalized to the corresponding holdout set, yielding 10000 independent at each sample size for each method. Such examined the true out of sample generalization performance for each method. The mean and 95% confidence intervals of the variance explained based on these 1000 samples is shown.

## RESULTS

### Simulation results

Figure 1 shows the predictive performance of the PVS-B, PVS-U, and Min-p on 2000 simulations of brain-behavior associations. The PVS-B consistently outperformed the PVS-U and the Min-p except at high signal sparsity level, and the differences were most evident at large sample sizes. The level of sparsity only impacted the predictive performance of the PVS-B at the most extreme condition, i.e. only 1% of the vertices contained true signal. At the sample size of the baseline ABCD data (*N* = 6103), the PVS-B showed 8 to 42 folds increase in the proportion of total signal explained compared to the Min-p when the true signal was not extremely sparse. No improvement in prediction was observed for the PVS-B compared to the Min-p when the true signal was restricted to 1% of the vertices.

**Figure 1.**
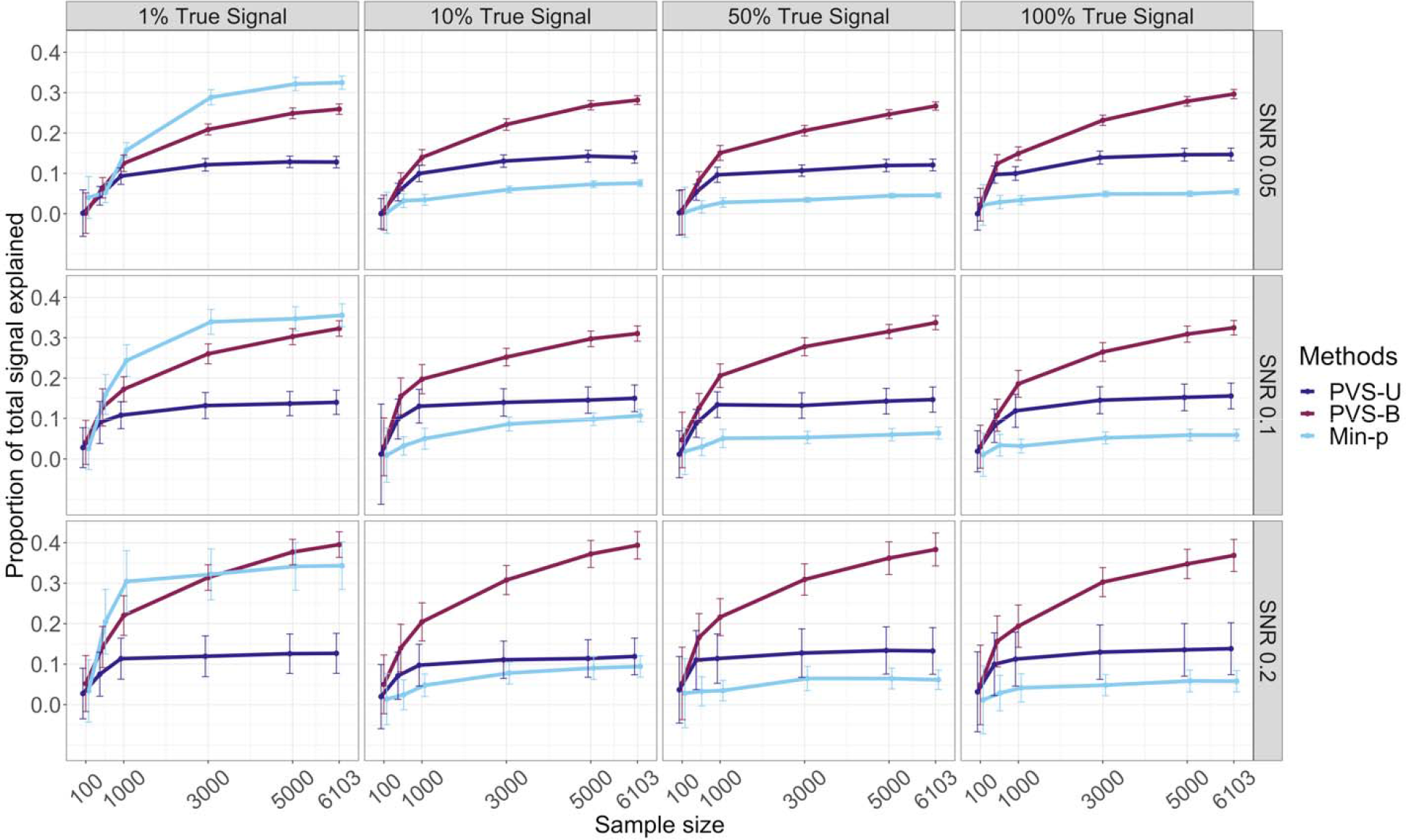
The predictive power of the PVS-U, PVS-B and Min-P using simulated data. The proportion of total signal explained, measured by the variance explained R^2^ of each method divided by the SNR level, for the PVS-U, PVS-B, and Min-p at varying simulated levels of proportion of true signal, SNR, and sample size. The PVS-B showed significantly greater variance explained than the PVS-U and Min-p except when the true signal was extremely sparse. Error bars show the 95% confidence interval of R^2^ across simulations.

We also evaluated how thresholding influenced the predictive performance of the PVS-B at varying levels of simulated proportion of true signal in Figure 2. The PVS-B demonstrated comparable, if not superior, prediction performance to thresholded PVS-Bs when the effect size per vertex was lower, comparable to many fMRI studies, (SNR of 0.05 and 0.1) and when the proportion of true signal was high (10%, 50% and 100%). The effect size per vertex was conceptualized as the SNR divided by the number of vertices with true signal. The unthresholded PVS-B was able to capture the predictive effect across the cortex when the true signal was global, however even at 10% sparsity level the unthresholded PVS-B performed equally well compared to thresholded versions at lower SNR. Only when the true signal was extremely sparse did the highly thresholded PVS-B perform better.

**Figure 2.**
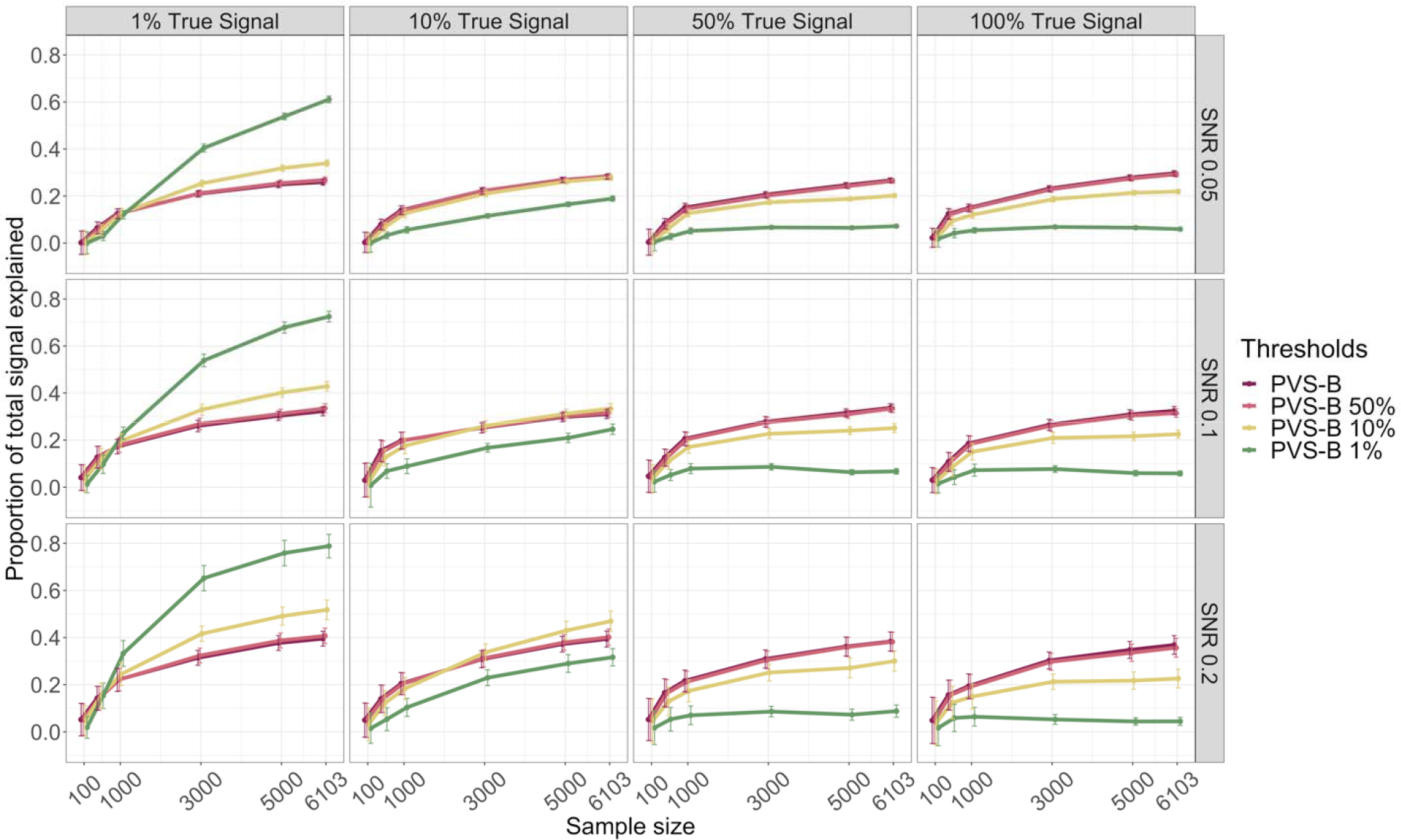
The predictive power of the PVS-B at different levels of thresholding. The proportion of total signal explained (R^2^) for each method divided by the SNR level, for the PVS-B and its thresholded variants at varying simulated levels of proportion of true signal, SNR, and sample size. 95% CIs were reported. The unthresholded PVS-B showed comparable to superior predictive performance compared to the thresholded PVS-Bs when the true signal was not extremely sparse.

### Empirical data results

We examined how well the PVS-U and PVS-B methods captured brain-behavior associations empirically by applying them to two fMRI task contrasts to predict individual differences in cognitive performance. To examine the sensitivity and specificity of the PVS predictive performance, four brain-behavior associations were investigated: nBack predicting TC, nBack predicting SSRT, SST predicting TC, and SST predicting SSRT. The prediction accuracy of the PVS-B, the PVS-U and Min-P for each brain-behavior association with 95% bootstrap CIs is reported in Table 1.

**Table 1.** Variance explained, with 95% bootstrap CI, for the four brain-behavior associations (empirical data) estimated for the PVS-B, the PVS-U, and the Min-p.

|  | PVS-B | PVS-U | Min-P |
| --- | --- | --- | --- |
| **nBack-TC** | 11.3<br>[9.8 - 12.9] | 7.5<br>[5.7 – 8.9] | 3.8<br>[2.7 – 4.8] |
| **nBack-SSRT** | 0.7<br>[0.2 - 1.1] | 0.2<br>[0 – 0.5] | 0.5<br>[0.1 – 0.8] |
| **SST-SSRT** | 6.9<br>[5.4 - 8.1] | 1.1<br>[0.6 – 1.5] | 0.4<br>[0.1 – 0.7] |
| **SST-TC** | 0.5<br>[0.1 - 0.8] | 0.2<br>[0 – 0.3] | 0.4<br>[0 – 0.6] |

For the nBack-TC association, the PVS-B demonstrated significantly better predictive performance compared to the PVS-U and the Min-p, explaining 11.3% of the individual variability in TC with the vertex-wise BOLD signal variation of the 2back-0back contrast of the nBack task. Similar improvement in prediction accuracy was also observed for the SST-SSRT association. The PVS-B was able to explain 6.9% of the variance in SSRT using the vertex-wise BOLD variation of the correct stop vs. correct go contrast from SST, compared to 1.1% for the PVS-U and 0.4% for the Min-p. For the nBack predicting SSRT and the SST predicting TC conditions, all methods showed minimal variance explained and the CIs bounded or were close to 0. The vertexwise BOLD signal variations in nBack and SST were not informative in explaining individual differences in SST and TC respectively.

We also examined how the predictive performance of the PVS-B varied as a function of thresholding for these brain-behavior associations (Figure 3). Thresholding based on the rank of the vertex-wise p-values resulted in decreased prediction accuracy for the two significant associations, suggesting that information from vertices with p-values that would not have reached vertex-wise corrected levels of statistical significance was still informative for behavioral prediction. Specifically, for the nBack-TC association, the PVS-B 50% and PVS-B 10% showed slightly reduced predictive performance (11.1% and 9.4%) compared to the PVS-B (11.3%), but the difference was not statistically significant. For the SST-SSRT association, the PVS-B, PVS-B 50% and PVS-B 10% yielded similar predictive performance, explaining 6.9%, 6.9% and 5.6% of the variance in SSRT using SST. These data show that using information from all or at least 50% if the vertices for prediction improves prediction accuracy, therefore it would be more appropriate to assume these associations are distributed across the cortex rather than sparse. For these two associations, all methods showed improvement in predictive performance with increased sample sizes (Supplementary Figure 2).

**Figure 3.**
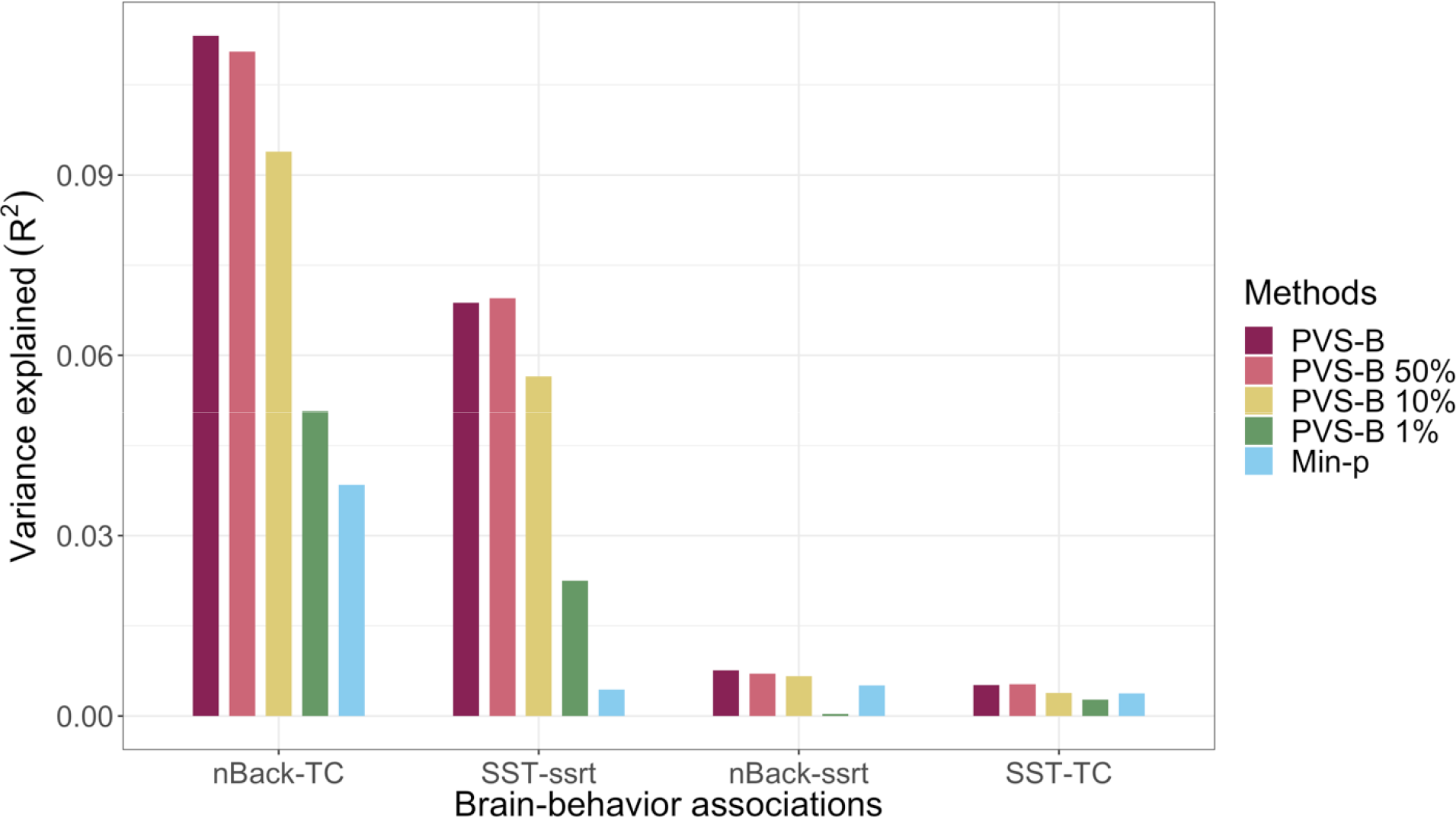
Predictive performance across PVS methods for the empirical brain-behavior associations. Variance explained for the four brain-behavior associations (nBack predicting TC, SST predicting SSRT, nBack predicting SSRT, SST predicting TC) estimated using PVS-B and its thresholded variants. For the two significant associations, the best prediction performance was achieved by the PVS-B where all vertices where included in the model. Predictive performance decreased as more stringent thresholds were applied.

## DISCUSSION

In this study, we have introduced the polyvertex score (PVS) framework. The PVS can be used to assess the magnitude of a brain-behavior association of interest by aggregating the explanatory power of neuroimaging data across the whole cortex to predict individual variability in behavior. The Bayesian form of the PVS incorporates the mass univariate parameter estimates of a brain-behavior association and the correlation structure of the imaging phenotype of interest to estimate the strength of the generalizable effect of the brain in an independent sample. Previous studies have shown that brain regions or vertices significantly contributing to behavioral variability each only account for a small proportion of behavioral variation (Poldrack et al., 2017; Smith & Nichols, 2017). Here we have shown that a whole brain phenotype, captured by the PVS-B, explained more individual variation in behavior compared to a subset of most significant vertices. With simulations, we have demonstrated that the PVS-B was able to handle varying signal structures and yielded optimal performance under varying levels of sample size, SNR, and proportion of true signal. Using the empirical data from the ABCD study, we also demonstrated that the PVS-B had superior predictive performance compared to the PVS-U and the Min-p, showing the importance of capturing the multivariate nature of the imaging phenotype for optimal prediction.

Traditional methods of analyzing neuroimaging data aim to identify brain regions significantly associated with a behavior of interest. This approach has been fruitful in characterizing the involvement of brain regions and systems in behavior, but has produced little utility in predicting behavioral variability. Using simulations and empirical data, we have shown that incorporating effect sizes across all vertices, using the PVS method, explained more variance in behavior than thresholding vertices based on significance level. Consistent with association patterns reported in other task-based fMRI studies (Chang et al., 2015; Gonzalez-Castillo et al., 2012) and for other brain phenotypes (Dubois et al., 2018; Palmer et al, in prep; Smith et al., 2015), the explanatory effect of these functional brain phenotypes on the behaviors tested was widespread and distributed across the cortex. Our results provide empirical evidence that it is important to account for the whole brain pattern of associations when studying brain-behavior relationships. Moreover, the PVS-B also demonstrated specificity in prediction such that significant relationships were identified specifically for the nBack-TC and the SST-SSRT association but not for the other two brain-behavior associations of interest. The PVS-B captures the available explanatory effect without overfitting to boost prediction performance.

Nevertheless, there are limitations to our current approach. First, the signal architecture across the cortex varies in sparsity and distribution depending on the brain-behavior association of interest. Complex behaviors, such as general cognitive performance, are associated with activation patterns or cortical architectures across widespread brain regions (Bruin, Denys, & Wingen, 2019; Reddan, Lindquist, & Wager, 2017). More focused contrasts or specialized tasks, such as finger tapping or visual perception, on the other hand, may demonstrate sparser and more localized effects across the cortex; therefore, we would hypothesize that in these cases the unthresholded PVS-B would not predict individual variability better than the thresholded PVS-Bs. The superior empirical predictive performance of the unthresholded PVS-B suggests that the underlying signal structure of the association between the task fMRI data analyzed and cognitive performance is not sparse. The PVS-B does not explicitly impose a sparsity assumption, such as the spike-and-slap prior (Mitchell & Beauchamp, 1988), yet still demonstrates robustness against varying signal sparsity levels of the effect size distribution, making it an appealing predictive tool when the true association structure in the brain is unknown.

Secondly, the PVS-B only captures additive effects across the cortex as its parameters are calculated based on the mass univariate parameter estimates. Therefore, non-linear effects and interactions between vertices are not accounted for with the PVS-B. While the PVS-B may potentially show inferior performance to multivariate prediction models that assess nonlinear relationships across vertices, it is tightly linked to the traditional brain mapping approach and has important utilities for smaller sample studies to boost power for prediction. The PVS-B can be applied to a smaller scale imaging study where the mass univariate parameter estimates and the correlation structure of the imaging phenotype are acquired from the ABCD study to generate a more accurate estimate of a brain-behavior association in this independent sample. These PVS-B scores can subsequently be associated with other brain and behavioral phenotypes, mirroring the utility of the PRS in assessing the relationship between individual risks for disease and variability in behavior in genetics.

The application of the PVS-B prediction framework is not limited to functional imaging phenotypes. The PVS-B has proven useful for structural imaging phenotypes to examine the association between regional cortical morphology and cognition in children (Palmer et al., in prep). It can be applied to data of other imaging modalities and resolutions. A common challenge of multimodal imaging studies is the growing dimension of the parameter space by including predictors from multiple imaging modalities. The PVS-B tackles this issue by summarizing the effect sizes at all vertices as one composite measure. Thus, the number of parameters for multimodal analyses is significantly reduced, and PVS-Bs of different imaging modalities can be combined to examine the effect of multiple imaging properties of cortical and subcortical regions on behavior. As the PRS empowered the genetics research community in data sharing and scientific discovery, we believe that the PVS framework will shed light on the multivariate nature of brain-behavior associations, and will hopefully inspire more data sharing and collaborative effort among neuroimaging studies for more accurate and replicable scientific discoveries.

## ACKNOWLEDGEMENT

Data used in the preparation of this article were obtained from the Adolescent Brain Cognitive Development (ABCD) Study (https://abcdstudy.org), held in the NIMH Data Archive (NDA). The ABCD Study is a multisite, longitudinal study designed to recruit more than 10,000 children age 9-10 and follow them over 10 years into early adulthood. It is supported by the National Institutes of Health and additional federal partners under award numbers U01DA041022, U01DA041028, U01DA041048, U01DA041089, U01DA041106, U01DA041117, U01DA041120, U01DA041134, U01DA041148, U01DA041156, U01DA041174, U24DA041123, U24DA041147, U01DA041093, and U01DA041025. A full list of supporters is available at https://abcdstudy.org/federal-partners.html. A listing of participating sites and a complete listing of the study investigators can be found at https://abcdstudy.org/Consortium_Members.pdf. ABCD consortium investigators designed and implemented the study and/or provided data but did not all necessarily participate in analysis or writing of this report. This manuscript reflects the views of the authors and may not reflect the opinions or views of the NIH or ABCD consortium investigators. The ABCD data repository grows and changes over time. The data was downloaded from the NIMH Data Archive ABCD Collection Release 2.0.0 (DOI: 10.15154/1503209).

## SUPPLEMENTARY MATERIALS

**Supplementary Figure 1.**
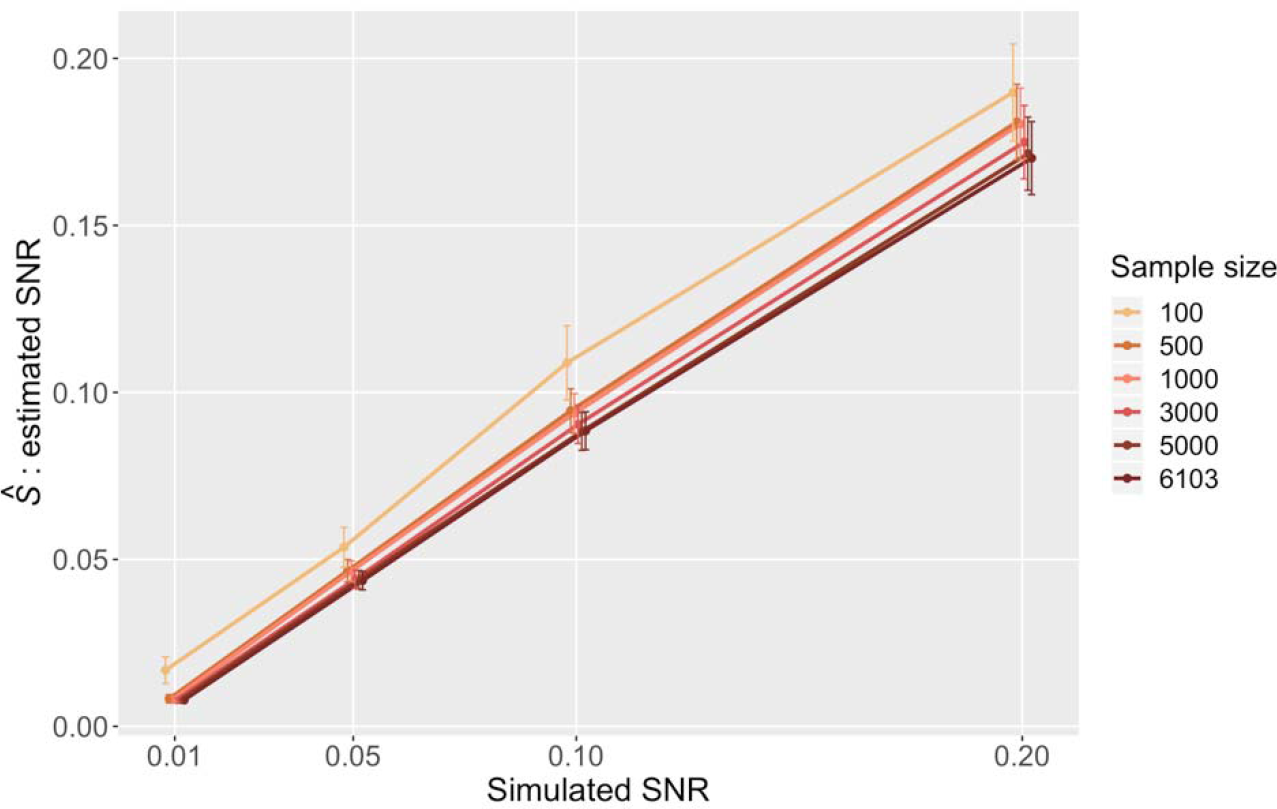
The estimated SNR at the corresponding true SNR level as a function of sample size in the simulations. The reliably estimated the true SNR across all sample sizes as demonstrated in simulations.

**Supplementary Figure 2.**
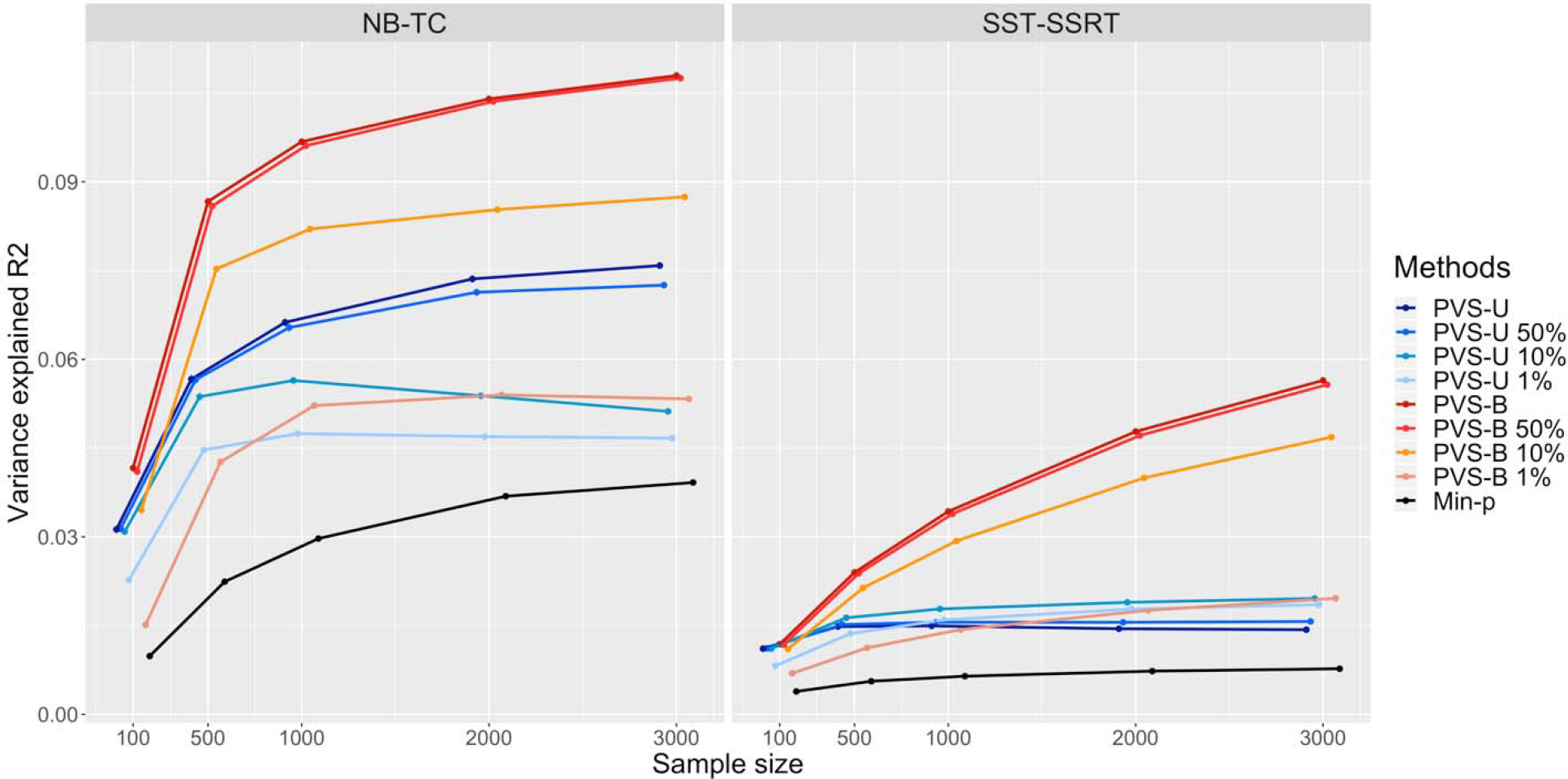
Variance explained and 95% CI based on 1000 subsamples of the full sample using each of the nine PVS methods as a function of sample size for the nBack predicting TC and the SST predicting SSRT condition.

